# Influenza A virus agnostic receptor tropism revealed using a novel biological system with terminal sialic acid-knockout cells

**DOI:** 10.1101/2022.02.21.481323

**Authors:** Haruhiko Kamiki, Shin Murakami, Takashi Nishikaze, Takahiro Hiono, Manabu Igarashi, Yuki Furuse, Hiromichi Matsugo, Hiroho Ishida, Misa Katayama, Wataru Sekine, Yasushi Muraki, Masateru Takahashi, Akiko Takenaka-Uema, Taisuke Horimoto

**Affiliations:** Laboratory of Veterinary Microbiology, Graduate School of Agricultural and Life Sciences, The University of Tokyo, Yayoi, Bunkyo-ku, Tokyo 113-8657, Japan; Koichi Tanaka Mass Spectrometry Research Laboratory, Shimadzu Corporation, Nishinokyo-Kuwabaracho, Nakagyo-ku, Kyoto 604-8511, Japan; Molecular and Cellular Glycoproteomics Research Group, Cellular and Molecular Biotechnology Research Institute, National Institute of Advanced Industrial Science & Technology, Tsukuba, Ibaraki 305-8565, Japan; Division of Global Epidemiology, International Institute for Zoonosis Control, Hokkaido University, Kita-ku, Sapporo 001-0020, Japan; Institute for Frontier Life and Medical Sciences, Kyoto University, Kyoto 606-8507, Japan; Iwate Medical University, Yahaba-cho, Iwate 028-3694, Japan; Research Institute for Environmental Sciences and Public Health of Iwate Prefecture, Morioka, Iwate 020-0857, Japan

## Abstract

Avian or human influenza A viruses bind preferentially to avian- or human-type sialic acid receptors, respectively, indicating that receptor tropism is an important factor for determining the viral host range. However, there are currently no reliable methods for analyzing receptor tropism biologically under physiological conditions. Here, we established a novel system using MDCK cells with avian- or human-type sialic acid receptors and with both sialic acid receptors knocked out (KO). When we examined the replication of human and avian influenza viruses in these KO cells, we observed unique viral receptor tropism that could not be detected using a conventional solid-phase sialylglycan binding assay, which directly assesses physical binding between the virus and sialic acids. Furthermore, we serially passaged an engineered avian-derived H4N5 influenza virus, whose PB2 gene was deleted, in avian-type receptor-KO cells stably expressing PB2 to select a mutant with enhanced replication in KO cells; however, its binding to human-type sialylglycan was undetectable using the solid-phase binding assay. These data indicate that a panel of sialic acid receptor-KO cells could be a useful tool for determining the biological receptor tropism of influenza A viruses. Moreover, the PB2-KO virus experimental system could help to safely and efficiently identify the mutations required for avian influenza viruses to adapt to human cells that could trigger a new influenza pandemic.

**Author summary:** Influenza A virus initiates infection via hemagglutinin by binding to avian- or human-type receptors. The acquisition of mutations that allow avian virus hemagglutinins (HAs) to recognize human-type receptors is mandatory for the transmission of avian influenza viruses to humans, which could lead to a pandemic. Therefore, it is important to detect such mutation(s) in animal influenza viruses for pandemic surveillance and risk assessment. In this study, we established a novel system using a set of genetically engineered MDCK cells with knocked out sialic acid receptors to biologically evaluate the receptor tropism for influenza A viruses. Using this system, we observed unique receptor tropism in several virus strains that was undetectable using conventional solid-phase binding assays that measure physical binding between the virus and artificially synthesized sialylglycans. This study makes a significant contribution to the literature because our findings suggest the pitfall of conventional receptor binding assay and the existence of a sialic acid-independent pathway for viral infection. In addition, our system could be safely used to identify mutations that could acquire human-type receptor tropism. Thus, this system could contribute not only toward basic analyses, such as elucidating the mechanism of influenza virus host range determination, but also the surveillance of viruses of animal origin that could be capable of infecting via human-type receptors, triggering a new influenza pandemic.

## Introduction

Influenza A virus initiates infection via viral hemagglutinin (HA) by binding to avian- or human-type receptors which consist of terminal sialic acids linked to galactose with α2,3 or α2,6 linkages, respectively [1, 2]. Human influenza viruses preferentially bind to human-type receptors, whereas avian influenza viruses bind to avian-type receptors. The human upper respiratory tract predominantly expresses human-type receptors, leading to the efficient infection and replication of human influenza viruses, but not avian influenza viruses [3, 4]. The acquisition of mutations on avian virus HAs to recognize human-type receptors is mandatory for the transmission of avian influenza viruses or reassortants with avian virus HAs to humans, which could lead to a pandemic. Therefore, it is important to detect such mutations in animal influenza viruses as genetic markers for surveillance and risk assessment for pandemic preparedness.

The binding affinity between HAs and sialic acid receptors is commonly measured using assays that employ synthesized short sialylglycans or enzymatically-treated red blood cells resembling human- or avian-type receptors [5–7]. Although these conventional methods are useful for measuring the physical binding affinity between HAs and sialic acids, they are unable to assess the biological correlation between sialylglycan binding and viral infection since binding affinity does not necessarily reflect the physiological state of sialylglycans on the cell surface, which display structural complexity in terms of their length and branching [8, 9]. In addition, different sialic acid forms are important for viral infectivity, such as N-acetylneuraminic (Neu5Ac)-type acid glycans, which act as functional receptors, and N-glycolylneuraminic acid (Neu5Gc)-type acid glycans, which act as pseudo-receptors to inhibit the infection of some influenza viruses [10].

Cells lacking sialic acids could be useful for assessing the preference of biological receptors for viral infection; however, no studies have yet reported any available cell lines lacking α2,3- and/or α2,6-linked sialylglycans that can support the multistep replication of influenza A viruses. Previous reports have indicated that SLC35A1 transports cytidine-5′-monophospho-N-acetylneuraminic (CMP-sialic) acid, a sialyltransferase substrate, from the cytoplasm to the Golgi apparatus [11], and that β-galactosidase α2,3 sialyltransferases (ST3Gals) or α2,6 sialyltransferases (ST6Gals) catalyze the transfer of CMP-sialic acids with α2,3 or α2,6 linkages to terminal galactose in the Golgi apparatus, respectively [12, 13]. Mammalian genomes encode six ST3Gal subfamilies (ST3Gal1-6) and two ST6Gal subfamilies (ST6Gal1 and ST6Gal2). Therefore, it is predicted that sialylglycan-knockout (KO) cells could be generated by knocking out the genes involved in sialylglycan synthesis.

In this study, we generated a series of cells lacking human- and/or avian-type receptors by knocking out six ST3Gals, two ST6Gals, or the SLC35A1 gene in MDCK cells. We then used these KO cells to biologically evaluate the receptor tropism of a panel of influenza viruses. By serially passaging a replication-incompetent avian virus lacking the PB2 gene, to limit biosafety concerns, in the avian-type receptor KO cells, we adapted the virus and selected mutants with enhanced replication in the KO cells. Through these experiments, we examined the receptor specificity of influenza viruses that have been overlooked in previous glycan-binding assays.

## Results

### Establishment of sialic acid-KO cells

To establish an assay system to biologically evaluate the receptor tropism of influenza viruses, we generated Madin-Darby canine kidney (MDCK) cells lacking human-and/or avian-type sialylglycans. First, we knocked out six ST3Gal (ST3Gal1-ST3Gal6) and/or two ST6Gal (ST6Gal1 and ST6Gal2) genes in the cells using a CRISPR/Cas9 system with guide RNAs targeting these genes (S1 Table). We also knocked out the sialic acid transporter SLC35A1 gene in these cells. After confirming in/del sequences in the target genomic DNA regions, we confirmed the absence of avian- or human-type sialylglycans on the KO cells by staining with anti-Neu5Acα2,3Gal monoclonal antibody (HYB4) and *Sambucus nigra* agglutinin (SNA) lectin recognizing Neu5Acα2,6Gal (Fig 1A). Wild-type (WT) MDCK cells were stained with both HYB4 antibody and SNA lectin, whereas ST3Gal-KO (human-type) cells were stained only with SNA lectin and ST6Gal-KO (avian-type) cells were stained only with HYB4 antibody. Both ST3Gal and ST6Gal-KO (DKO) cells and SLC35A1-KO (SLC35A1KO) cells were stained with neither SNA lectin nor HYB4 antibody. Together, these data indicate that human- and/or avian-type receptor KO cells were successfully established.

**Fig 1.**
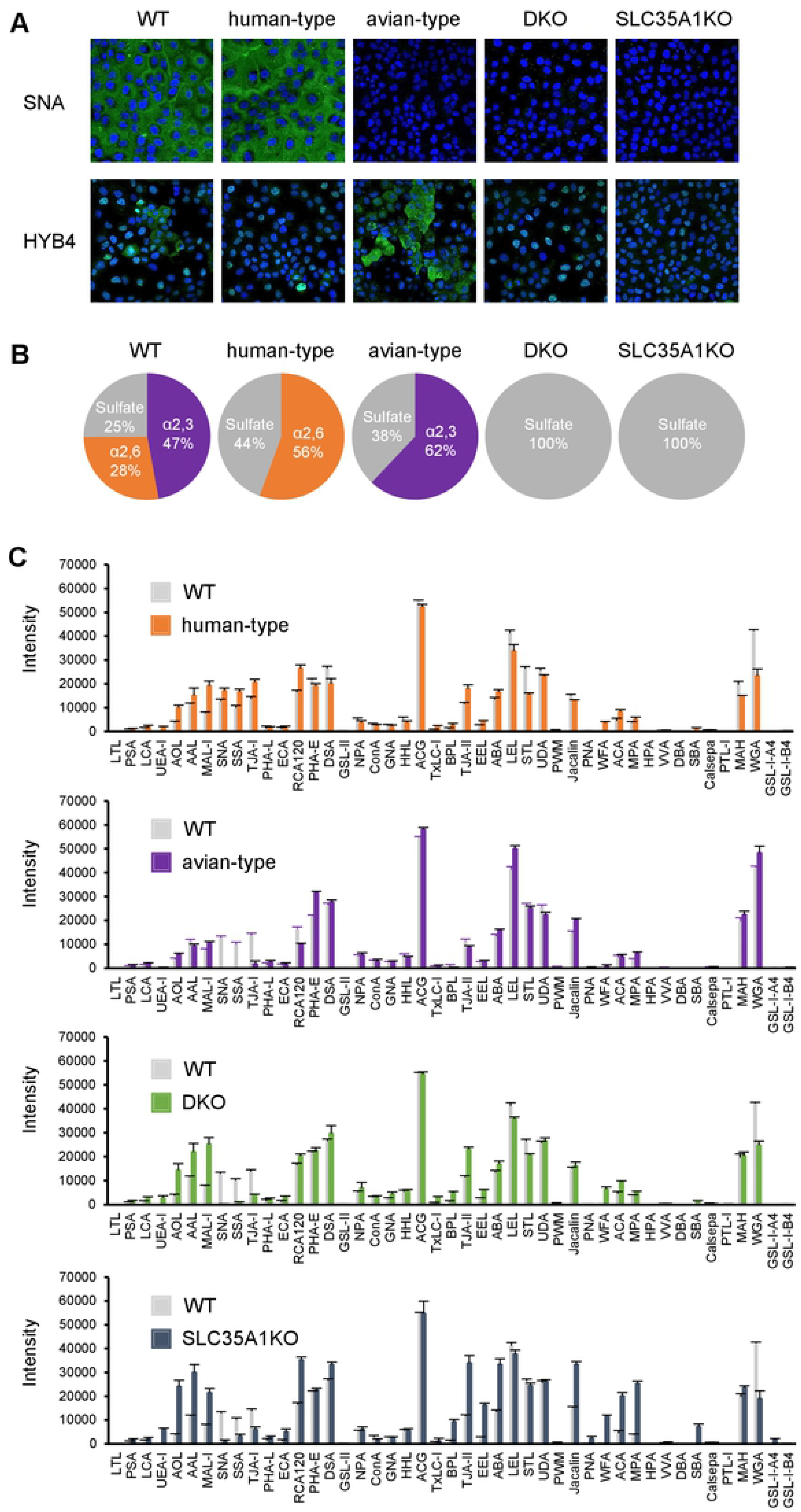
Confirmation of the presence of avian- and human-type sialylglycans in KO cells. (A) Wild-type (WT) MDCK and KO cells were stained with SNA, a lectin that recognizes Neu5Acα2,6Gal, and HYB4, a monoclonal antibody that recognizes Neu5Acα2,3Gal. Cell nuclei were counterstained with Hoechst 33342. (B) Acidic sialylglycans among the N-type glycans were detected using mass-spectrometric analysis with the SALSA method. The percentages of avian-type (α2,3), human-type (α2,6), and sulfated glycans are presented in pie charts in purple, orange, and gray, respectively. (C) The glycan profiles of WT, human-type, avian-type, DKO, and SLC35A1KO cells were analyzed using lectin arrays. Cells were biotinylated and lysed, and the proteins in the lysate were added to slides printed with 45 types of lectin. The amounts of proteins binding to the lectins were measured using a GlycoStation Reader 1200. Data were mean-normalized and presented as the mean of three replicates ± standard deviation.

### Mass-spectrometry and lectin array analyses of KO cells

Influenza viruses predominantly use N-linked sialylglycans, but not O-linked sialylglycans, during infection [14]. To investigate the distribution of N-linked glycans in KO cells, they were collected from the cell lysates and analyzed by mass spectrometry using the sialic acid linkage-specific alkylamidation (SALSA) method. This is based on the principal that α2,6- and α2,3-linked sialic acids are specifically amidated with different length alkyl chains that can be distinguished using mass spectrometry [15, 16]. High-mannose-type glycans were abundant in WT cells and all tested KO cells, while neutral glycans were abundant and acidic glycans were less abundant in DKO and SLC35A1KO cells than in WT, human-type, and avian-type cells (S1 Fig). We further analyzed acidic glycans, which include sialylglycans (Fig 1B), and found that while WT cells had both α2,3- and α2,6-linked sialylglycans, α2,3-linked glycans were more dominant. Human- and avian-type cells lacked α2,3- and α2,6-linked sialylglycans, respectively, whereas DKO and SLC35A1KO cells lacked sialylglycans. The levels of terminal-sulfated galactose increased in all KO cells, while all acidic glycans consisted of terminal sulfated galactose in DKO and SLC35A1KO cells (Fig 1B).

Next, we investigated the glycan profiles of KO cells using lectin microarray assays [17, 18]. A panel of lectins were immobilized as arrays (S2 Table) and incubated with fluorescently labeled cell lysates containing all N- and O-linked glycans. The interactions between the lectins and lysates were detected using a fluorescence imager. Signals for MAL-I lectin, which recognizes Siaα2,3Gal and sulfated galactose [19], were higher in human-type, DKO, and SLC35A1KO cells than in WT cells (Fig 1C). Since Siaα2,3Gal was completely absent in these cells, as indicated by mass-spectrometry analysis (Fig 1B), these increased signals should be due to sulfated galactose, consistent with the mass-spectrometry results (S2 Fig). The signals for the Siaα2,6Gal-recognizing lectins, SNA and SSA, and the Siaα2,6Galβ1,4GlcNAc-recognizing lectin, TJA-I, were also higher in human-type cells than in WT cells, indicating an increase in the amount of α2,6-linked sialic acid due to ST3Gal gene knock out. Conversely, SNA, SSA, and TJA-I signals were substantially lower in avian-type, DKO, and SLC35A1KO cells (Fig 1C), further supporting the mass spectrometry results. These results, in combination with those of the mass-spectrometry analysis, suggest that the KO cells lacked N-linked α2,3- and/or α2,6-linked sialic acid residues and likely corresponding O-linked glycans.

### Viral growth in sialic acid-KO cells and physical binding to avian- and human-type synthetic sialylglycans

Each avian H4N5, H6N8, or H10N4 virus grew efficiently in both WT and avian-type cells but displayed limited growth in human-type, DKO, and SLC35A1KO cells. These findings indicate that the avian viruses preferentially use avian-type receptors for their growth, consistent with their specific physical binding to avian-type sialylglycans in the solid-phase glycan binding assay (Fig 2A). Osaka/H1N1pdm09 and its descendant Iwate/2019 grew efficiently not only in WT and human-type cells, but also in avian-type cells, albeit at a slightly lower rate for Osaka/H1N1pdm09. However, the growth of both viruses was greatly reduced in both DKO and SLC35A1KO cells, indicating that these human H1N1 viruses use both human- and avian-type receptors for growth, despite their specific physical binding to human-type sialylglycan in the glycan binding assay (Fig 2B). The recent seasonal Iwate/H3N2 virus only grew well in human-type cells, suggesting that it has strict human-type receptor selectivity for growth, although it displayed unclear physical binding to sialylglycans in the glycan binding assay (Fig 2B). The human-derived laboratory strain PR8 (H1N1) grew efficiently in WT and avian-type cells but much less in human-type cells (Fig 2C), suggesting that successive passages of this human virus in chicken embryonated eggs might lead to preferential binding specificity to avian-type receptors, as supported by a strong physical binding to avian-type sialylglycans (Fig 2C). Another human-derived laboratory strain Aichi/68(H3N2), which is the ancestor of the seasonal H3N2 virus with unknown passage history, grew efficiently in both human- and avian-type cells, which did not correlate with the physical binding outcome of the glycan binding assay (Fig 2C). The avian-origin HK/99(H9N2) virus, which was isolated from humans, is known to have a high affinity for human-type sialylglycan as it possesses HA with L at position 226 [20, 21]. This was consistent with the findings of our glycan binding assay and the observation that HK/99(H9N2) grew efficiently in both human- and avian-type cells as well as WT cells (Fig 2D). Taken together, these results suggest that the physical binding profiles between the viruses and sialic acids provided by the solid-phase glycan binding assay do not necessarily correlate with the biological usage of the receptors for viral growth.

**Fig 2.**
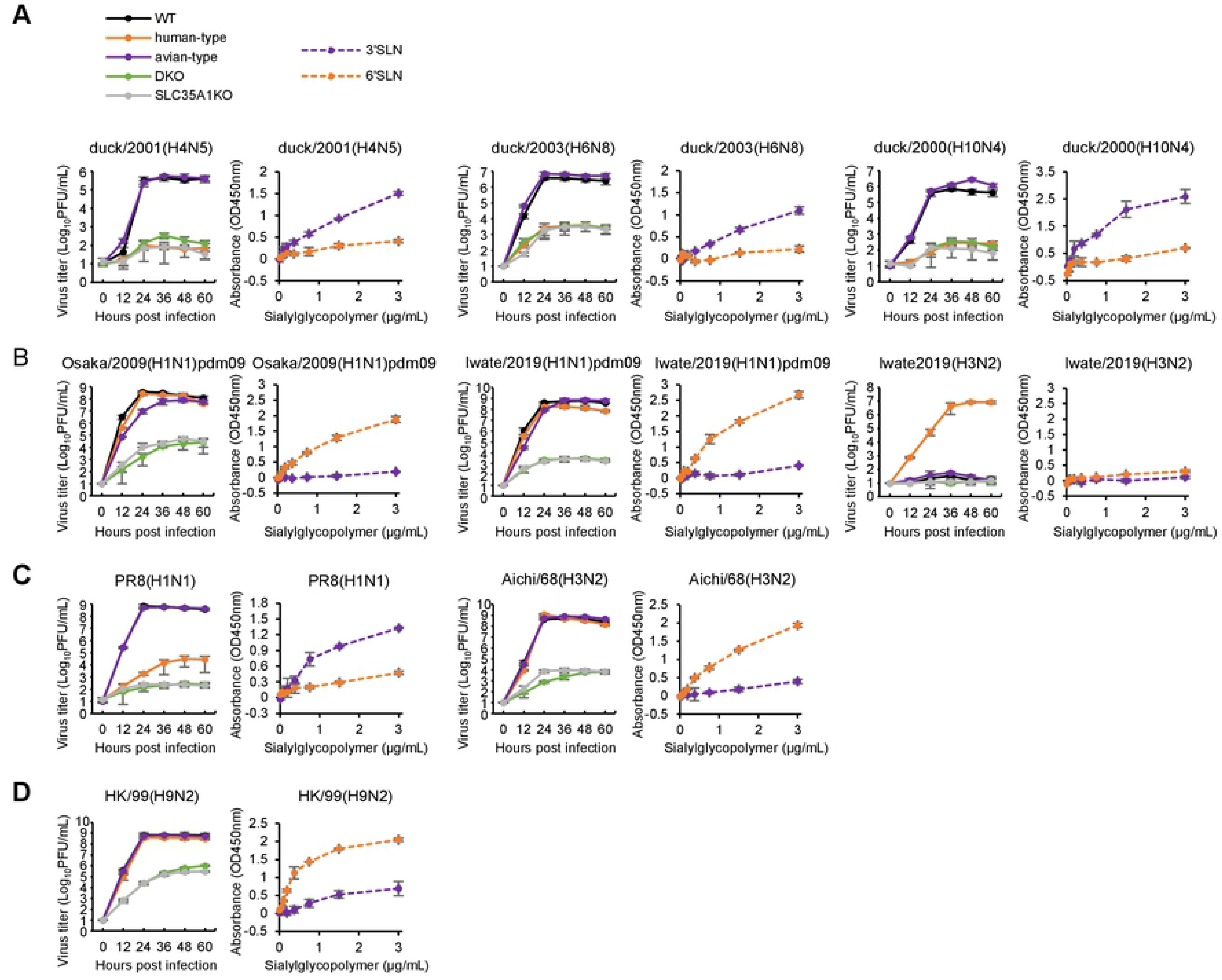
Comparison of virus growth in sialic acid-KO cells and their physical binding to sialylglycopolymers. Wild-type (WT) and sialic acid-KO cells were infected with viruses at an moi of 0.01. Supernatants were collected every 12 h and viral titers were measured using plaque assays. Error bars indicate the standard deviation of the mean of three independent experiments. Virus growth in WT, human-type, avian-type, DKO, and SLC35A1KO cells is indicated by black, orange, purple, green, and gray solid lines, respectively. For the solid-phase binding assay, the virus (64 HAU) was absorbed to ELISA plate which was blocked and reacted with biotinylated avian-type (3’SLN) or human-type (6’SLN) sialylglycopolymers, followed by HRP-labeled biotin-streptavidin complex. TMB substrate reagent was added to develop color and the reaction was stopped using 2% sulfuric acid. Absorbance was measured at 450 nm. Data represent the mean of three independent experiments ± standard deviation. The binding properties of the virus to avian- and human-type sialylglycopolymers are shown using purple and orange dashed lines, respectively. The growth and physical sialylglycan binding of (A) avian-derived, (B) human-derived, (C) laboratory, and (D) H9N2 viruses are shown on the left and right, respectively. The area under the curve (AUC) was calculated to compare growth curves among samples. Statistical significance of the differences for AUC between each type and WT cells was analyzed using ANOVA with post hoc Dunnett’s multiple comparisons test and shown in *p* value (S5 Table).

### Virus growth in sialic acid-addback cells

To confirm that the viral growth properties in KO cells (Fig 2) were not due to off-target mutations from the CRISPR/Cas9 system, we generated four types of sialic acid-addback cells in which the ST3Gal1 gene, ST6Gal1 gene, both genes, or the SLC35A1 gene were reintroduced into human-type, avian-type, DKO, or SLC35A1KO cells, respectively. Sialic acid expression was restored in each addback cell type, as confirmed by staining with SNA lectin and HYB4 antibody (S3 Fig). We then tested viral growth in these cells, finding that growth was restored to comparable levels as in the parental KO cells (S4 Fig). These results indicate a lack of off-target mutations and suggest that the difference in viral growth between the WT and KO cells was caused by KO of the corresponding sialylglycan.

### Adaptation of an avian virus to human-type cells

Although it is important to identify mutations that increase viral growth in human cells in order to monitor avian influenza viruses and assess their pandemic potential [22, 23], it is difficult to predict these mutations under laboratory conditions. Therefore, we used our established human-type cells to produce an evaluation system. We used reverse genetics to generate a replication-incompetent PB2-knockout virus [24] possessing HA and NA segments of the avian H4N5 virus, in addition to a modified PB2 segment containing the green fluorescent protein (GFP) gene instead of the PB2 gene and the remaining five segments from WSN (namely rH4/PB2KO virus). We also established WT, human-, and avian-type cells stably expressing PB2 from WSN, which were termed WT/PB2, human-type/PB2, and avian-type/PB2 cells, respectively. We serially passaged the rH4/PB2KO virus in human-type/PB2 cell lines independently six times (lines A to F). Although the parental virus did not show clear cytopathic effects (CPEs) in the cells, they became clear by the sixth passage in all virus lines. Sequencing indicated that five out of the six passaged virus lines possessed Q to L or R mutations at amino acid position 226 of HA (HA1-Q226L/R), whereas line C possessed HA1-I111L and HA2-A44S without an HA1-226 mutation (Table 1). Line F had HA2-D112A as well as HA1-Q226R and NA-G341R, although HA2-A112 and NA-R341 were partially observed as a mixed population in the parental virus. HA1-226 and HA1-111 were located on the head domain of HA, and HA1-226 directly contacts the terminal sialic acid of the glycan receptor. HA2-44 and HA2-112 were located on the stalk region of HA (S5 Fig).

**Table 1.** Mutations detected in human-type/PB2 cell-adapted PB2-deficient H4N5 virus.

| Virus strain | HA | NA |
| --- | --- | --- |
| H4N5-P6-A | HA1-Q226L | — |
| H4N5-P6-B | HA1-Q226R | — |
| H4N5-P6-C | HA1-I111L<br>HA2-A44S | — |
| H4N5-P6-D | HA1-Q226R | — |
| H4N5-P6-E | HA1-Q226L | — |
| H4N5-P6-F | HA1-Q226R<br>HA2-D112A | G341R |

—, no mutation was detected.

### Growth of H4N5 mutant viruses in sialic acid-KO cells and binding to synthetic glycans

To identify the mutation(s) in the adapted rH4/PB2KO virus responsible for efficient growth in human-type/PB2 cells, we generated recombinant rH4/PB2KO viruses with HA1-Q226L, HA1-Q226R, HA1-I111L, and/or HA2-A44S, which were referred to as rH4/PB2KO-Q226L, rH4/PB2KO-Q226R, rH4/PB2KO-I111L, rH4/PB2KO-A44S, and rH4/PB2KO-I111L+A44S, respectively. The rH4/PB2KO virus grew efficiently in both WT/PB2 and avian-type/PB2 cells, but not in human-type/PB2 cells, and bound specifically to avian-type glycan (Fig 3A), as did the duck/2001(H4N5) virus (Fig 2A). The rH4/PB2KO-I111L, rH4/PB2KO-A44S, and rH4/PB2KO-I111L+A44S viruses grew efficiently in WT/PB2 and avian-type/PB2 cells, like the rH4/PB2KO virus, but grew more than the parental virus in human-type/PB2 cells (Fig 3B, 3E and 3F). These mutant and parental viruses bound specifically to avian-type glycan in the glycan-binding assay, consistent with the viral growth observed in KO cells; however, the differences in growth between the mutant and parental strains could not be explained by the glycan-binding assay. rH4/PB2KO-Q226L grew more efficiently in human-type/PB2 than in WT/PB cells, but grew at similar levels in avian-type/PB2 cells and in WT/PB cells (Fig 3C), despite its specific binding to human-type glycans. By contrast, rH4/PB2KO-Q226R grew efficiently in all cells, despite its specific binding to avian-type glycans (Fig 3D). These results confirm that the solid-phase glycan binding assay could not determine the biological usage of glycan receptors for viral growth and suggest that the KO cell-based virus adaptation assay could identify mutations that allow avian viruses to acquire efficient growth in human-type cells, unlike the glycan-binding assay.

**Fig 3.**
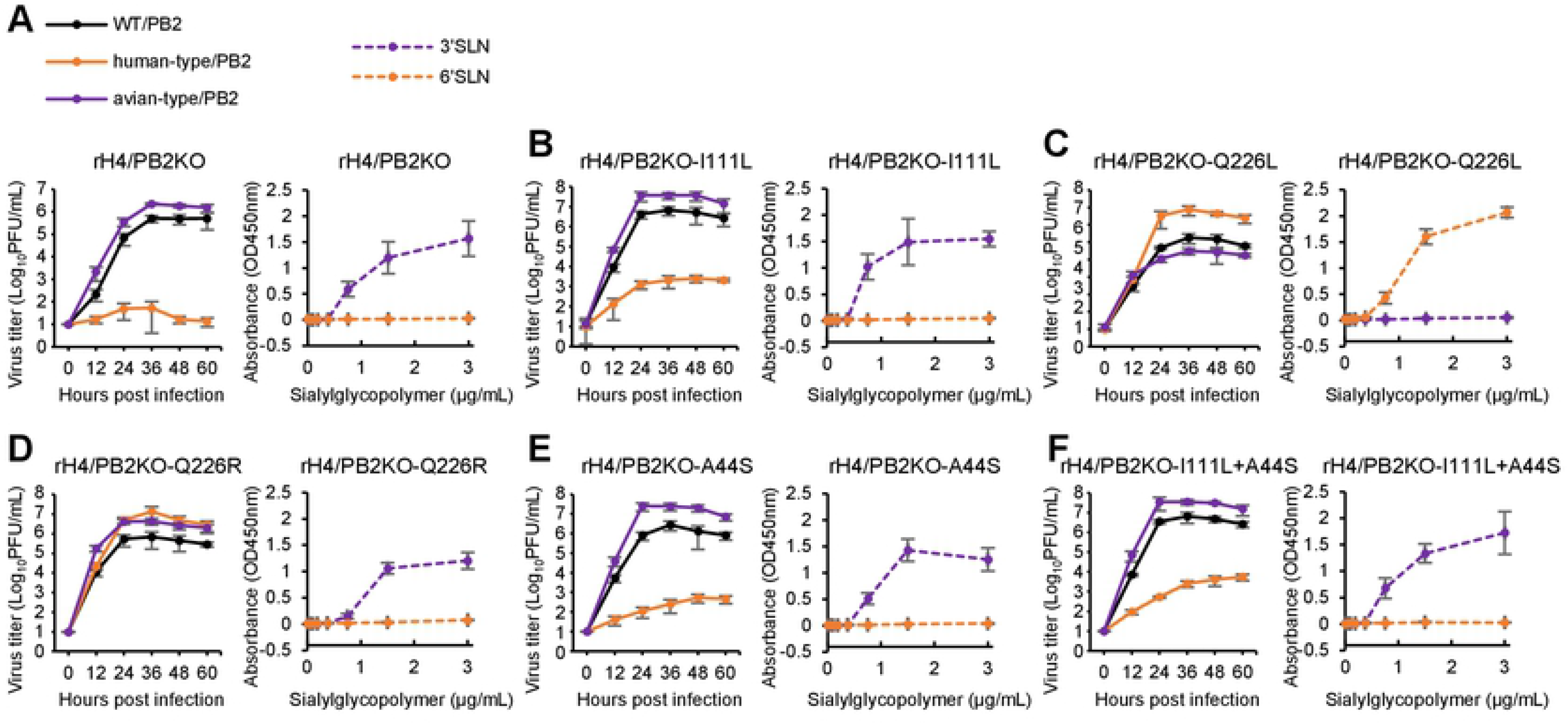
PB2-deficient H4N5 virus growth in PB2-expressing KO cells and binding activity to sialylglycopolymers. PB2-deficient viruses bearing wild-type (WT) or mutant H4 HA and N5 NA were inoculated into WT, human-type, and avian-type cells expressing PB2 at an moi of 0.01. Supernatants were collected every 12 h and the viral titer was measured using a plaque assay. Error bars indicate the standard deviation of the mean of three independent experiments. Virus growth in WT, human-type, and avian-type cells are indicated by black, orange, and purple solid lines, respectively. For the solid-phase binding assay, various amounts of sialylglycopolymers were absorbed to the ELISA plate. After blocking, viruses (32 HAU) were added to the plate and reacted with mouse antiserum against the H4N5 virus, followed by HRP-labeled secondary antibodies. TMB substrate reagent was added to develop color and the reaction was stopped using 2% sulfuric acid. Absorbance was measured at 450 nm. Data represent the mean of three independent experiments ± standard deviation. The binding properties of the virus to avian- and human-type sialylglycopolymers are shown as purple and orange dashed lines, respectively. The growth and physical binding to sialylglycans of (A) rH4/PB2KO, (B) rH4/PB2KO-I111L, (C) rH4/PB2KO-Q226L, (D) rH4/PB2KO-Q226R, (E) rH4/PB2KO-A44S, and (F) rH4/PB2KO-I111L+A44S virus are shown on the left and right, respectively. The area under the curve (AUC) was calculated to compare growth curves among samples. Statistical significance of the differences for AUC between each type/PB2 and WT/PB2 cells was analyzed using ANOVA with post hoc Dunnett’s multiple comparisons test and shown in *p* value (S6 Table).

### Molecular dynamics simulation between HAs and human-type glycan

Finally, we analyzed the molecular basis of the HA1-Q226L/R mutations during binding to the human-type sialyl pentasaccharides analog (LSTc) using molecular dynamics simulation. To evaluate the stability of the HA-LSTc complex, the root mean square deviation (RMSD), which is an index of deviation from the initial structure, was calculated for each HA. The binding free energies between LSTc and H4 HAs were also calculated using the MM-GBSA method. In HA1-226Q, the RMSD of the LSTc increased with time and dissociated from HA, suggesting weak LSTc binding, whereas in HA-HA1-226L or -226R, the RMSD did not change significantly over time, suggesting stable binding (Fig 4A). The binding energies of the LSTc to HA1-226Q, -226L, and -226R were -43.2, -47.7, and -47.3 kcal/mol, respectively, indicating that 226L/R binds to human-type glycans more strongly than 226Q. The interaction between HA and the LSTc over 60% of the simulation time was found to be at only one site for HA1-226Q, but at five and four sites for HA1-226L and HA1-226R, respectively (Fig 4B). Together, these results suggest that the HA1-Q226L/R mutation enhances binding to human-type sialylglycans.

**Fig 4.**
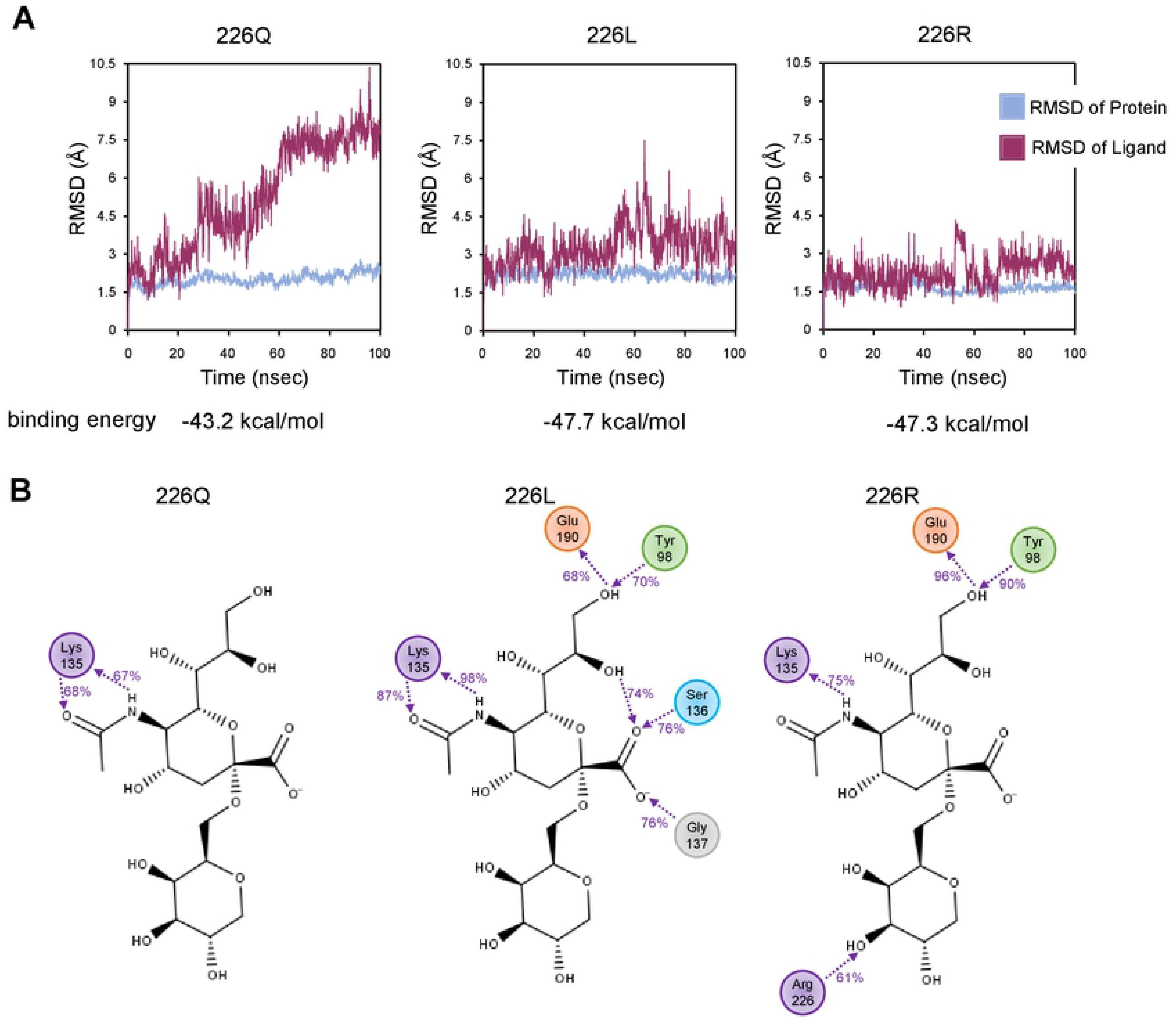
Molecular dynamics simulation of human-type glycan and HA protein from H4 subtype. (A) Plots of RMSD against simulation time (100 ns). Blue and red lines indicate the RMSD of HA proteins and the glycan analog LSTc (Neu5Acα2,6Galβ1,4GlcNAcβ1,3Galβ1,4Glc), respectively. Simulation results from HA protein bearing Q, L, or R, at amino acid position 226 in HA1 are shown. (B) Detailed interactions between LSTc and amino acid residues. Interactions between galactose/sialic acid and the HA protein that were detected over 60% of the simulation time are indicated. Numbers between the interacting molecules represent the ratio (%) of the time when the interaction was observed against the total simulation time.

## Discussion

The receptor tropism of influenza A viruses is one of the most important factors that determines their host range. To elucidate the mechanism via which avian viruses cross species barriers to infect humans and become pandemic viruses, it is essential to analyze their receptor specificity. Until now, receptor tropism analyses have mainly been performed by evaluating physical binding affinity to short sialylglycans that resemble avian-type α2,3- or human-type α2,6-linked sialylglycans. However, these assay systems are often technical, requiring skills and expertise, and the resulting physical binding profiles do not necessarily represent the biological binding of the virus to receptors for growth. Here, we developed a system that directly evaluates the biological receptor-tropism of viruses by detecting their replication in artificially manipulated cells with avian and/or human-type receptor KO. In addition, we developed a method to identify amino acid mutations in avian viruses that allow human-type receptor-mediated infection by adapting a replication-incompetent PB2-deficient virus with avian-derived HA to KO cells stably expressing PB2.

In this study, we found considerable inconsistency between the growth properties of some viruses in sialic acid-KO cells and their sialylglycan-binding affinities as determined using conventional solid-phase binding assays. The growth properties of the avian H4, H6, and H10 viruses correlated well with the results of the binding assay, whereas H1N1pdm2009, Aichi/68 (H3N2), and HK/99 (H9N2) grew efficiently in avian-type cells despite their low binding affinities to avian-type sialylglycans. As reported in recent studies [25, 26], it was difficult to detect the physical binding of recent seasonal H3N2 viruses, such as Iwate/2019, to human-type sialylglycans using the binding assays; however, this virus grew efficiently in human-type cells with human-type receptor preference [27]. These results indicate that our sialic acid-KO cell-based assay system can be used to assess the receptor specificity of viruses, even if their binding to avian- or human-type sialylglycans is undetectable using the binding assay. Thus, our assay system could be useful for biologically evaluating viral receptor preference. The acquisition of genetic mutations that increase binding affinity to human-type receptors is an essential factor for the adaptation of animal-derived viruses to humans. Passage experiments using primary human airway epithelial cells could be used to search for such mutations in avian viruses [28], since most cell lines like MDCK cells cannot be utilized as they possess both receptor types, despite supporting the efficient growth of most influenza A viruses [29]. These types of passage experiment can also present biosafety concerns as they generate mutant viruses with human-type receptor specificity. Therefore, we sought to create a replication-incompetent system using a PB2-deficient virus [24], wherein infectious viruses would not be produced in any intact cells. By combining this system with sialic acid-KO cells, we identified HA1-Q226L and HA1-Q226R mutations in the avian H4 virus that are important for recognizing the human-type receptors. HA1-Q226L is a key mutation for avian H3N2 and H5N1 viruses to acquire binding to human-type receptors [30, 31], and also contributes toward H5N1 and N7N9 viruses becoming transmissible between ferrets, which may indicate a potential for human-to-human transmissibility [22,23,32]. HA1-226R has been detected in human and swine H1N1, avian H5N1, and egg-adapted H1N1pdm viruses [33] (S4 Table); however, the effects of this residue varied among viruses. In the human H1N1pdm virus, HA1-Q226R increased binding to avian-type sialylglycan and decreased binding to human-type sialylglycan [33], whereas in avian H5 virus, this mutation did not change binding affinity to human-type sialylglycan but decreased binding to avian-type sialylglycan [34]. In this study, HA1-Q226R enhanced the preference of avian H4 virus for human-type sialylglycans, as evaluated using molecular dynamics simulations, but it did not change its binding affinity for human-type sialylglycan in the binding assay. Nonetheless, mutations at position 226 in HA1 may play a pivotal role in allowing avian viruses to change their biological binding to sialic acid-receptors and thus potentially infect humans. Such human infection may also trigger further mutations for adaptation during the course of replication in humans.

It is interesting to note that, even in terminal sialic acid-knockout DKO and SLC35A1KO cells, most viruses were able to grow, albeit less efficiently. These findings suggest the presence of a sialic acid receptor-independent pathway for viral infection and imply that an unidentified molecule(s) besides sialic acids may act as a functional receptor for viral infection. Previous studies have shown that other molecules, such as C-type lectins (DC-SIGN and L-SIGN) and MHC-class II, can bind the influenza virus to the cell surface [35]; however, it has not yet been determined whether these molecules act as biologically functional receptors for viral infection. We believe that sialic acid-KO cells such as those generated in this study could help to elucidate the molecular mechanism of early viral infection, including alternative receptors.

In conclusion, we developed a novel method to evaluate the receptor- preference of influenza viruses using viral replication as a biological indicator. In addition, we demonstrated that this method can be utilized to safely assess mutations for the adaptation of animal viruses to human cells that could lead to pandemic potential. This strategy could be used not only for basic analyses, such as elucidating the mechanism of viral host range determination, but also for the surveillance of viruses of animal origin that are capable of infection via human-type receptors.

## Materials and methods

### Cells and viruses

Madin-Darby canine kidney (MDCK) cells were obtained from the American Type Culture Collection (ATCC; CCL-34) and maintained in minimal-essential medium (MEM) supplemented with 5% newborn calf serum (NCS) and antibiotics. Human embryonic kidney 293T cells were obtained from the RIKEN BioResource Research Center (RCB2202) and maintained in Dulbecco’s modified Eagle’s medium (DMEM) supplemented with 10% fetal bovine serum (FBS) and antibiotics. All cells were incubated in 5% CO_2_ at 37 °C.

The A/duck/Hokkaido/1058/01 (H4N5), A/duck/Hokkaido/228/03 (H6N8), A/duck/Hokkaido/18/2000 (H10N4), A/Osaka/364/2009 (H1N1), A/Iwate/34/2019 (H1N1), A/Puerto Rico/8/34 (H1N1), and A/Aichi/2/68 (H3N2) viruses were propagated in embryonated chicken eggs or MDCK cells. The A/Iwate/43/2019 (H3N2) virus was propagated in avian-type receptor-KO MDCK cells, which were established in this study. The A/Hong Kong/1073/99 (H9N2) virus was generated using reverse genetics [36] and propagated in MDCK cells. Propagated viruses were aliquoted and stored at -80 °C.

### Generation of sialic acid-KO cells

Avian-type receptor-KO (human-type) cells, human-type receptor-KO (avian-type) cells, avian- and human-type receptor-KO (DKO) cells, and SLC35A1-KO (SLC35A1A1KO) cells were generated by knocking out the corresponding genes using a CRISPR/Cas9 system. The target sequences for six ST3Gals (ST3Gal-1, -2, -3, -4, -5, and -6), two ST6Gals (ST6Gal-1 and -2), and the SLC35A1 gene were designed using CRISPR direct (https://crispr.dbcls.jp/; S1 Table) and each sequence was cloned into plentiCRISPR plasmids [37] (Addgene plasmid #52961, a gift from Dr. Feng Zhang) using NEBuilder (New England Biolabs, Ipswich, MA, USA). MDCK cells were transfected with the six ST3Gal-targeting plasmids to generate human-type cells, the two ST6Gal-targeting plasmids to generate avian-type cells, or the SLC35A1-targeting plasmid to generate SLC35A1KO cells using polyethylenimine (PEI; Polysciences, Warrington, PA, USA). At 24 h post-transfection, the cell supernatant was replaced with medium containing 5 μg/mL puromycin. Drug-resistant clones were randomly selected and their genomic DNA was sequenced. Cells possessing in/dels in all targeted genes were chosen for further analysis. DKO cells were generated by transfecting human-type cells with ST6Gal-targeting plasmids, followed by the same process as the other KO cells.

### Addback of genes involved in glycan sialylation to KO cells

RNA was extracted from MDCK cells using ISOGEN (NIPPON Genetics, Tokyo, Japan), and cDNA was prepared by reverse transcription with oligo dT and RevaTraAce (TOYOBO, Osaka, Japan). The coding sequences of ST3Gal1, ST6Gal1, and SLC35A1 were amplified using PCR with specific primers and cloned into the pS lentiviral vector [38] using NEBuilder (New England Biolabs).

To prepare recombinant lentivirus vectors expressing ST3Gal1, ST6Gal1, or SLC35A1, pS plasmids were co-transfected with p8.9QV [38] and pCAGGS-VSV-G, which express vesicular stomatitis virus G protein in 293T cells using PEI. At 48 h post-transfection, recombinant lentiviruses in the cell supernatant were concentrated using polyethylene glycol (PEG) precipitation and inoculated into the corresponding KO cells for gene addback. Human-type cells with ST3Gal1 addback were referred to as human-type/ST3Gal1 cells. Avian-type/ST6Gal1, DKO/ST3,6Gal1, and SLC35A1KO/SLC35A1 cells were also generated.

### Lectin and antibody staining of KO and addback cells

To stain the KO and addback cells, we used lectin *Sambucus nigra* agglutinin (SNA, biotinylated: Vector Laboratories, Burlingame, CA, USA) that recognizes Neu5Acα2,6Gal and the antibody HYB4 [39] (FUJIFILM, Tokyo, Japan) that recognizes Neu5Acα2,3Gal. Cells were plated on an 8-well chamber slide (AGC Techno Glass, Shizuoka, Japan), fixed with 4% paraformaldehyde for 5 mins, and blocked with skimmed milk at 21 °C for 1 h. To stain Neu5Acα2,3Gal glycans, SNA (10 μL/mL in PBS) was added to the cells at 150 μL/well, incubated at 21 °C for 1 h, washed with PBS, and incubated with DyLight488 conjugate streptavidin (Vector Laboratories) (10 μL/mL in PBS) at 150 μL/well at 21 °C for 1 h. To stain Neu5Acα2,3Gal glycans, HYB4 (15 μL/mL in PBS) was added to the cells at 150 μL/well, incubated at 4 °C overnight, washed with PBS, and incubated with goat anti-mouse IgG conjugated to Alexa 488 (Abcam, Cambridge, UK; 1:400 dilution in PBS) at 150 μL/well at 21 °C for 1 h. Cell nuclei were counterstained with Hoechst 33342 (Dojindo, Kumamoto, Japan). Stained cells were observed under a confocal microscope (Carl Zeiss LSM700; ZEISS, Oberkochen, Germany).

### Mass spectrometry analysis of N-linked glycans using the SALSA method

N-linked glycans were analyzed using mass spectrometry according to the sialic acid linkage-specific alkylamidation (SALSA) method, as described previously [15, 16]. α2,3 and α2,6-linked sialic acids were specifically amidated with different length alkyl chains twice by alkylamidation, resulting in distinct linkage with different molecular masses. We prepared the following reagents for sialic acid derivatization: 1-(3-(dimethylamino)propyl)-3-ethylcarbodiimide hydrochloride (EDC-HCl), 1-hydroxybenzotriazole monohydrate (HOBt), isopropylamine hydrochloride (iPA-HCl), dimethyl sulfoxide (DMSO), methanol (MeOH), and aqueous methylamine (MA(aq)) (Tokyo Chemical Industry, Tokyo, Japan). After the wild-type MDCK, human-type, avian-type, DKO, and SLC35A1KO cells had been washed with PBS, they were lysed in Pierce^TM^ IP Lysis Buffer (Thermo, Waltham, MA, USA) and centrifuged to remove debris. Glycoproteins in the lysates were denatured using sodium dodecyl sulfate (SDS). After cooling on ice, N-linked glycans were released using PNGase F and directly applied to hydrazide beads (BlotGlyco^TM^ kit; Sumitomo Bakelite, Tokyo, Japan). The beads were added to the first alkylamidation solution (500 mM EDC-HCl, 500 mM HOBt, 2 M iPA-HCl in DMSO) and incubated at 25 °C for 1 h with gentle agitation. The beads were washed three times with MeOH, before being added to the second alkylamidation solution (10% MA(aq)) and gently agitated for a few seconds. After three washes with water, derivatized N-glycans were released from the beads in their native forms and their reducing ends labeled with 2-aminobenoic acid (2AA). N-glycans were ionized with 2,5-dihydroxybenzoic acid (DHB) and mass spectra were obtained using MALDI-QIT-TOF-MS (AXIMA-Resonance, Shimadzu/Kratos, Manchester, UK) and MALDI-TOF-TOF-MS (MALDI-7090, Shimadzu/Kratos) in negative ion mode.

### Lectin array analysis of glycoproteins from wild-type and sialic acid-KO cells

Lectin microarray analysis was performed as described previously with some modifications [17, 18]. Briefly, wild-type and sialic acid-KO MDCK cells were seeded on the 60 cm dish. The cells were washed with ice-cold PBS for three times and labeled with 1 mg/mL EZ-Link Sulfo-NHS-Biotin (Thermo Fisher Scientific) at 4 °C for 1 h. The cells were then washed with TBS to quench excess biotin-labeling reagents and lysed with TBS containing 1% Nonidet P-40 alternative, 0.5% sodium deoxycholate, 0.1% SDS, and protease inhibitor (Complete Ultra Mini; Roche, Basel, Switzerland). Proteins were then diluted with probing buffer and applied to an activated LecChip (ver.1.0; GlycoTechnica, Kanagawa, Japan; S2 Table). The array glass slide was incubated at 20°C overnight. After washing, the slides were overlayed with Streptavidin, Alexa Fluor 555 Conjugate (Thermo Fisher Scientific), and scanned using a GlycoStation Reader 1200 (GlycoTechnica). The obtained images were processed using GlycoStation ToolsPro Suite (ver.1.5). The net intensity of each lectin was calculated from the mean value of three spots minus the background.

### Generation of PB2-stably expressing cells

PB2-stably expressing wild-type MDCK, human-type, and avian-type cells were generated (WT/PB2, human-type/PB2, and avian-type/PB2 cells, respectively). The PB2 coding sequence from A/WSN/33(H1N1) was cloned into the pCAGGS-Blast plasmid [40], which was transfected into wild-type, human-type, and avian-type cells with PEI. The transfected cells were treated with 10 μg/mL of blasticidin (Kaken Pharmaceutical, Tokyo, Japan) and drug-resistant cells were randomly cloned. PB2 expression was confirmed by observing GFP-expressing PB2-deficient virus propagation [24].

### Evaluation of virus growth in cells

Cells were inoculated with viruses at a multiplicity of infection (moi) of 0.01 and maintained in MEM containing 0.3% bovine serum albumin (MEM/BSA) and 1 μg/mL of L-(tosylamido-2-phenyl) ethyl chloromethyl ketone (TPCK)-trypsin (Worthington, Lakewood, NJ, USA). Supernatants were collected every 12 h and viral titers were measured using a plaque assay.

### Generation of PB2-KO H4N5 virus

The HA and NA segments of the duck/2001(H4N5) virus were cloned into the RNA-expression plasmid pHH21 [41]. Wild-type and mutant PB2-KO H4N5 viruses were generated using reverse genetics, as described previously [41]. Briefly, wild-type or mutant HA and NA from duck/2001(H4N5), PB1, PA, NP, M, and NS from A/WSN/33, and the pPolIPB2(300)GFP(300) viral RNA-expressing plasmid [42] were transfected together with PB2, PB1, PA, and NP from A/WSN/33 protein-expressing plasmids into 293T cells. At 48 h post-transfection, supernatants containing PB2-KO viruses were harvested, propagated in WT/PB2 cells, and stored at -80 °C.

### Adaptation of H4N5 virus to human-type/PB2 cells

Six dishes (6 cm diameter) containing human-type/PB2 cells were infected with the PB2-KO H4N5 and maintained in MEM/BSA with 1 μg/mL TPCK trypsin. Three days post-infection, viruses were collected from the supernatants and inoculated into fresh human-type/PB2 cells. After six passages, the viruses were collected and their entire genome was Sanger-sequenced.

### Evaluation of virus binding to synthetic glycans

Solid-phase binding assays were performed as described previously [43], with some modifications. Briefly, viruses were pelleted by ultracentrifugation at 2500 rpm for 3 h using a P32ST rotor (CP80WX; Himac, Tokyo, Japan) through a 30% sucrose cushion. The pellets were suspended in a small volume of PBS and the protein concentration as HA titer were measured. Each virus was diluted in PBS to 64 HA/100 μL (ranging from 2.2 to 9.1 μg/mL of protein per virus; Supplementary Table 3) and added (100 μL/well) to ELISA plates (Maxisorp Nunc-immuno plates; Thermo). Since Iwate/2019(H3N2) did not show any detectable HA titer, as reported previously [22], the virus was diluted with a protein concentration of 5.1 μg/mL in PBS per well. Once the viruses had adsorbed to the wells at 4 °C overnight, they were washed with ice-cold PBS, 4% BSA in PBS was added (200 μL/well), and the plate was blocked at 21 °C for 2 h. After two washes with ice-cold PBS, the wells were reacted with 50 μL of 3’SLN (Neu5Acα2,3Galβ1,4GlcNAcβ) or 6’SLN (Neu5Acα2,6Galβ1,4GlcNAcβ) artificial synthetic glycans conjugated with biotin (3’SLN-C3-BP or 6’SLN-C3-BP, respectively; GlycoNZ, Auckland, New Zealand) diluted to 3, 1.5, 0. 75, 0.375, 0.1875, 0.09375, and 0.046875 μg/mL in reaction buffer (RB; 0.02% BSA, 1 μM oseltamivir (Wako Chemicals, Japan) in PBS) at 4 °C overnight. The wells were then washed twice with PBS and reacted with avidin/biotin-HRP complex (VECTASTAIN Elite ABC kit; Vector Laboratories) at 50 μL/well at 4 °C for 1 h. After a further five washes, 100 μL 3,3’,5,5’-tetramethyl-benzidene (TMB) substrate (Vector Laboratories) was added to each well and incubated at 21°C for 12 min to detect a reaction. Next, the reaction was stopped with 2% sulfuric acid (25 μL/well), and absorbance was measured at 450 nm using a microplate reader (iMark; Bio-Rad, Tokyo, Japan).

Solid-phase binding assays were used to reduce non-specific reactions to assess PB2-KO H4N5 viruses. Briefly, ELISA plates were incubated with streptavidin (100 μL/well, diluted to 1 μg/100 μL in PBS) at 21 °C for 3 h [44], washed twice with PBS, and incubated overnight at 4 °C with artificial synthetic glycopolymers (3’SLN-C3-BP or 6’SLN-C3-BP; GlycoNZ), diluted in PBS to 3, 1.5, 0.75, 0.375, 0.1875, 0.09375, 0.09375, and 0.046875 μg/mL at 50 μL/well. After blocking with Pierce Protein-Free (TBS) Blocking Buffer (200 μL/well; Thermo Fisher Scientific) at 21 °C for 3 h, the plates were incubated overnight at 4 °C with 100 μL of H4 PB2-deficient viruses (rH4/PB2KO, -I111L, -Q226L, -Q226R, - A44S, or -I111L+A44S) diluted to 32 HA/100 μL in PBS. The plates were then washed twice with ice-cold PBS, reacted with mouse antiserum against duck/2000(H4N5) (1:100 in dilution buffer; Can Get Signal solution 1; TOYOBO) at 50 μL/well at 4 °C for 1 h, washed a further two times, and incubated with anti-mouse IgG secondary antibodies with HRP (#NA9340, GE Healthcare) (1:4000) at 50 μL/well at 4 °C for 1 h. After a further five washes, the plates were reacted with TMB substrate (100 μL/well; Vector Laboratories) at 21 °C for 12 min and 2% sulfuric acid was added (25 μL/well) to stop the reaction. The absorbance was measured at 450 nm using a microplate reader.

### Molecular dynamics simulation of human-type glycan and HA protein

The three-dimensional (3D) structure of H4-HA, generated by X-ray crystallography with 2.10 Å resolution, was downloaded from the Protein Data Bank archive (PDB ID: 5XL7) to provide data for HA possessing HA1-226L combined with human-type glycan analog LSTc (Neu5Acα2,6Galβ1,4GlcNAcβ1,3Galβ1,4Glc). The mutagenesis wizard tool (PyMOL software) was used to generate mutant HA possessing Q or R at position 226 in HA1. Molecular dynamics simulation was conducted using the Desmond simulation package [45] (Schrödinger Release 2019–1: Desmond Molecular Dynamics System: DE Shaw Research: New York, NY, 2019). Based on the position of HA at the beginning of the simulation, the RMSD of the protein at each time point was calculated for 100 ns. The RMSD of the LSTc was calculated based on its relative position to HA. The binding free energies of the LSTc and HA were evaluated using the MM-GBSA method [46].

## Acknowledgments

We thank Dr. Yoshihiro Sakoda (Hokkaido University) for providing the viruses.

